# Multiple pathways for SARS-CoV-2 resistance to nirmatrelvir

**DOI:** 10.1101/2022.08.07.499047

**Authors:** Sho Iketani, Hiroshi Mohri, Bruce Culbertson, Seo Jung Hong, Yinkai Duan, Maria I. Luck, Medini K. Annavajhala, Yicheng Guo, Zizhang Sheng, Anne-Catrin Uhlemann, Stephen P. Goff, Yosef Sabo, Haitao Yang, Alejandro Chavez, David D. Ho

**Affiliations:** Aaron Diamond AIDS Research Center, Columbia University Vagelos College of Physicians and Surgeons, New York, NY, USA; Division of Infectious Diseases, Department of Medicine, Columbia University Vagelos College of Physicians and Surgeons, New York, NY, USA; Integrated Program in Cellular, Molecular, and Biomedical Studies, Columbia University Vagelos College of Physicians and Surgeons, New York, NY, USA; Medical Scientist Training Program, Columbia University Vagelos College of Physicians and Surgeons, New York, NY, USA; Department of Pathology and Cell Biology, Columbia University Vagelos College of Physicians and Surgeons, New York, NY, USA; Shanghai Institute for Advanced Immunochemical Studies and School of Life Science and Technology, ShanghaiTech University, Shanghai, China; Department of Microbiology and Immunology, Columbia University Vagelos College of Physicians and Surgeons, New York, NY, USA; Department of Biochemistry and Molecular Biophysics, Columbia University Vagelos College of Physicians and Surgeons, New York, NY, USA

## Abstract

Nirmatrelvir, an oral antiviral targeting the 3CL protease of SARS-CoV-2, has been demonstrated to be clinically useful in reducing hospitalization or death due to COVID-19^1,2^. However, as SARS-CoV-2 has evolved to become resistant to other therapeutic modalities^3–9^, there is a concern that the same could occur for nirmatrelvir. Here, we have examined this possibility by *in vitro* passaging of SARS-CoV-2 in increasing concentrations of nirmatrelvir using two independent approaches, including one on a large scale in 480 wells. Indeed, highly resistant viruses emerged from both, and their sequences revealed a multitude of 3CL protease mutations. In the experiment done at a larger scale with many replicates, 53 independent viral lineages were selected with mutations observed at 23 different residues of the enzyme. Yet, several common mutational pathways to nirmatrelvir resistance were preferred, with a majority of the viruses descending from T21I, P252L, or T304I as precursor mutations. Construction and analysis of 13 recombinant SARS-CoV-2 clones, each containing a unique mutation or a combination of mutations showed that the above precursor mutations only mediated low-level resistance, whereas greater resistance required accumulation of additional mutations. E166V mutation conferred the strongest resistance (~100-fold), but this mutation resulted in a loss of viral replicative fitness that was restored by compensatory changes such as L50F and T21I. Structural explanations are discussed for some of the mutations that are proximal to the drug-binding site, as well as cross-resistance or lack thereof to ensitrelvir, another clinically important 3CL protease inhibitor. Our findings indicate that SARS-CoV-2 resistance to nirmatrelvir does readily arise via multiple pathways *in vitro*, and the specific mutations observed herein form a strong foundation from which to study the mechanism of resistance in detail and to inform the design of next generation protease inhibitors.

## Main text

The COVID-19 (coronavirus disease 2019) pandemic has continued to affect the global populace. The rapid development and deployment of effective vaccines as well as monoclonal antibody therapeutics beginning in late 2020 have helped to greatly curtail its impacts^10–16^. Yet, the etiologic agent, SARS-CoV-2 (severe acute respiratory syndrome coronavirus 2), has continuously evolved to develop resistance to antibody-mediated neutralization^4–8^. In particular, several of the recent Omicron subvariants exhibit such strong antibody resistance that vaccines have had their protection against infection dampened and a majority of current monoclonal therapeutics have lost efficacy^4,5,8^, as manifested by increasing breakthrough infections in convalescing and/or vaccinated individuals^3^.

Fortunately, treatment options remain. In the United States, three antivirals have received emergency use authorization for COVID-19 treatment: remdesivir^17,18^, molnupiravir^19–21^, and nirmatrelvir^1,2^ (also known as PF-07321332, used in combination with ritonavir and marketed as PAXLOVID™). The first two target the RNA-dependent RNA polymerase (RdRp), and the latter targets the 3CL protease (3CL^pro^; also known as main protease (M^pro^) and nonstructural protein 5 (nsp5)). Both enzymes are essential for the viral life cycle and relatively conserved among coronaviruses^22,23^. Remdesivir is administered intravenously and has a reported relative risk reduction of 87%^18^, whereas molnupiravir and nirmatrelvir are administered orally and have reported clinical efficacies of 31%^20^ and 89%^1^, respectively, in lowering hospitalization or death. As the use of these antivirals increases, there is a concern that drug resistance may arise, particularly if given as monotherapies. For remdesivir, *in vitro* and *in vivo* studies have revealed mutations associated with resistance^9,24,25^, and resistance to molnupiravir or nirmatrelvir is now under active investigation. Here, we report that there are multiple routes by which SARS-CoV-2 can gain resistance to nirmatrelvir *in vitro*.

### Nirmatrelvir resistance in Vero E6 cells

To select for resistance to nirmatrelvir, SARS-CoV-2 (USA-WA1/2020 strain) was passaged in the presence of increasing concentrations of the drug (see **Methods** for details). We conducted this initial experiment in triplicate, using Vero E6 cells as they have been one of the standard cell lines used in COVID-19 research. After 30 passages, each of the three lineages demonstrated a high level of resistance, with IC_50_ values increasing 33- to 50-fold relative to that of the original virus (**Figs. 1a-d**). Examination of earlier viral passages confirmed a stepwise increase in nirmatrelvir resistance with successive passaging (**Figs. 1b-d**), without any evidence of resistance to remdesivir (**Fig. 1e**). The resistant viruses selected by passaging maintained their replicative fitness *in vitro*, with similar growth kinetics as those passaged without nirmatrelvir (**Extended Data Fig. 1**).

**Fig. 1.**
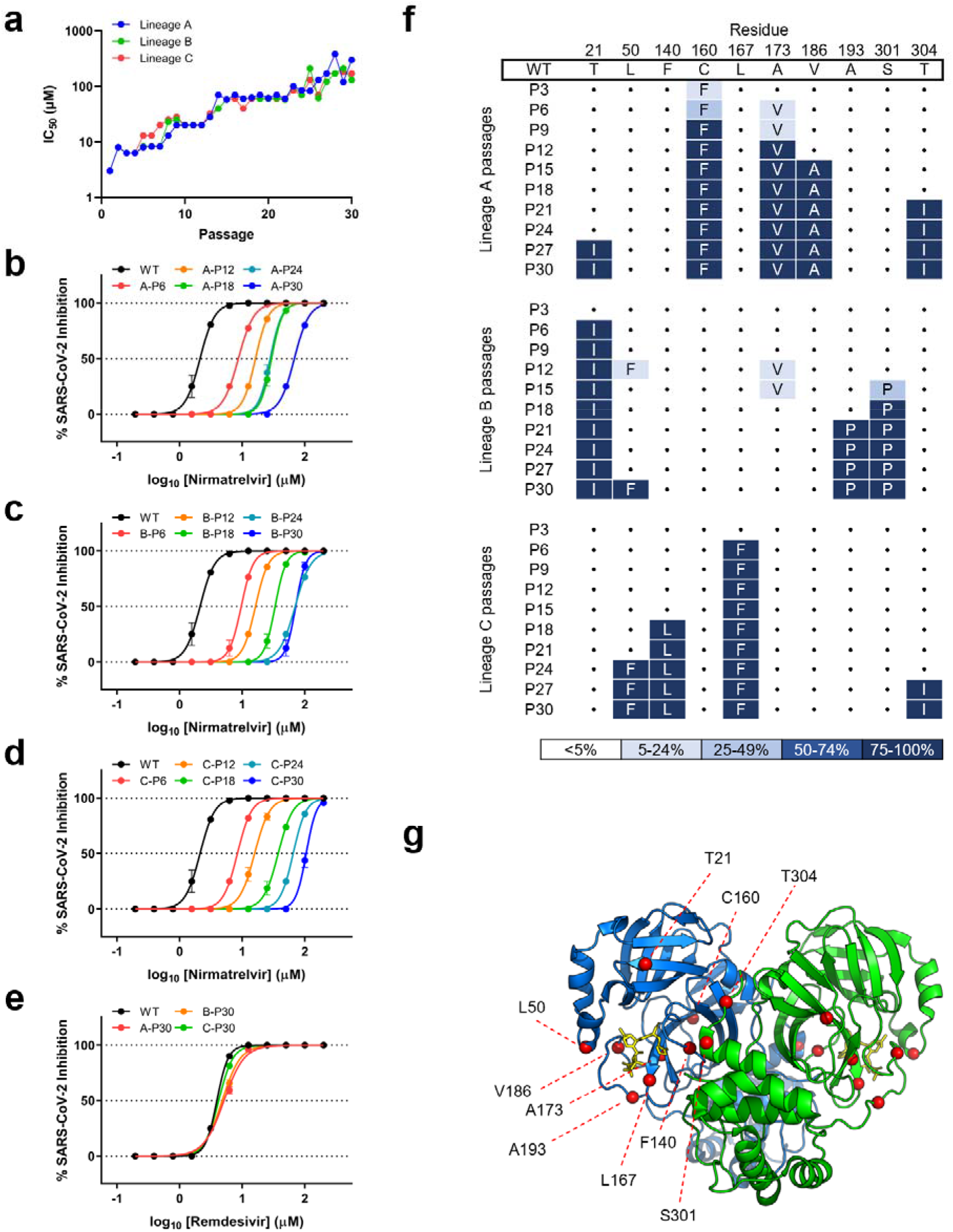
Identification of nirmatrelvir resistance in Vero E6 cells. **a,** Changes in IC_50_ during passaging of SARS-CoV-2 with nirmatrelvir. Vero E6 cells were infected in triplicate with SARS-CoV-2 (USA-WA1/2020) and passaged to fresh cells every 3 days for 30 passages. See Methods for additional details. **b-d,** Validation of nirmatrelvir resistance from the indicated passage from each of the three lineages, A, B, and C, respectively. **e,** Inhibition of passage 30 viruses from each lineage by remdesivir. **f,** Mutations in 3CL^pro^ found in the indicated passages from each lineage. Dots indicate wild-type at that residue. Mutations are shaded according to frequency. **g,** Residues mutated in passaging in Vero E6 cells overlaid onto the 3CL^pro^ structure with nirmatrelvir bound. The Cα of each residue that was mutated is denoted with a red sphere. The 3CL^pro^-nirmatrelvir complex was downloaded from PDB under accession code 7VH8. Error bars denote mean ± s.e.m of four technical replicates in **a-e**.

We then sequenced the 3CL^pro^ gene from the three viral lineages collected every three passages to investigate which mutations may confer resistance (**Fig. 1f**). We found that the three lineages harbored unique mutations, with only one mutation, at most, overlapping between the different lineages (T21I in lineages A and B, L50F in lineages B and C, and T304I in lineages A and C). The observed mutations occurred in a stepwise manner, mirroring the increases in drug resistance (**Fig. 1f**), and a number of them, but not all, were situated near the nirmatrelvir-binding site (**Fig. 1g**). Specifically, F140L and L167F were within 5 Å from nirmatrelvir. These results suggested that SARS-CoV-2 could readily develop nirmatrelvir resistance using several mutational pathways.

### Nirmatrelvir resistance in Huh7-ACE2 cells at scale

We therefore set out to conduct another passaging experiment to select for nirmatrelvir resistance, but this time at a larger scale with many replicates to better capture the multitude of solutions that SARS-CoV-2 could adopt under drug pressure. For these later studies, we utilized Huh7-ACE2 cells to examine if differences would arise in human cells, and because Vero E6 cells express high levels of P-glycoprotein, an efflux transporter that limits the intracellular accumulation of nirmatrelvir^26^. We passaged SARS-CoV-2-mNeonGreen (USA-WA1/2020 background with ORF7 replaced with mNeonGreen^27^) independently in 480 wells for 16 passages, with increasing concentrations of nirmatrelvir over time, and viruses from every fourth passage were subjected to next-generation sequencing (NGS) (**Fig. 2a** and see **Methods** for details). After 16 passages, varying degrees of nirmatrelvir resistance were observed as exemplified by the three viruses shown (**Fig. 2b**). Sequencing of the 3CL^pro^ in all wells that retained mNeonGreen signal identified 53 mutant populations (**Fig. 2c**). Across all of these populations, mutations were observed at 23 residues within the enzyme (1-6 mutations in each isolate), both proximal (≤ 5 Å; S144A, E166(A/V), H172(Q/Y), and R188G) and distal (> 5 Å) to nirmatrelvir (**Fig. 2d**). While there was widespread diversity among the passaged populations, seven mutations appeared ten or more times across replicates: T21I, L50F, S144A, E166V, A173V, P252L, and T304I. Mutations were only rarely observed across 3CL^pro^ cut sites within the polyproteins, except those at the termini of 3CL^pro^ itself (note that all but one example were in the C-terminal cut site), suggesting that substrate cleavage site alterations are largely not responsible for the nirmatrelvir resistance (**Extended Data Fig. 2**).

**Fig. 2.**
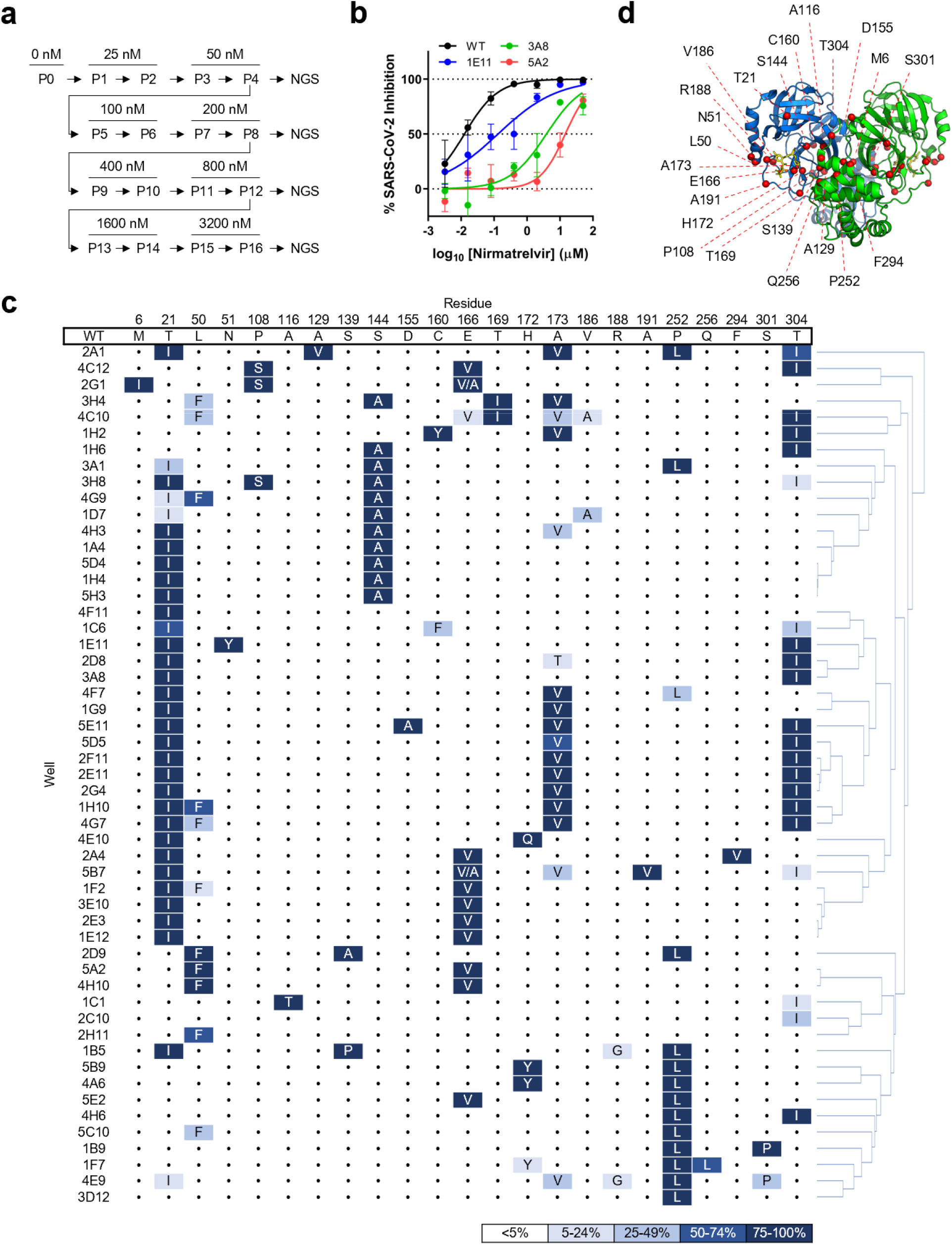
Identification of nirmatrelvir resistance at scale in Huh7-ACE2 cells. **a,** Passaging scheme. 480 wells were infected with SARS-CoV-2-mNeonGreen and passaged to fresh Huh7-ACE2 cells every 3-4 days, with the concentration of drug doubled every two passages. **b,** Validation of nirmatrelvir resistance of three wells from passage 16. These viral populations had the following mutations: 3A8 (T21I, T304I), 1E11 (T21I, N51Y, T304I), 5A2 (L50F, E166V). See **Supplemental Table 1** for exact frequencies. Representative curves from a single experiment from two biologically independent experiments are shown. Error bars denote mean ± s.e.m of three technical replicates. **c,** Mutations in 3CL^pro^ found in passage 16 from 53 wells. Dots indicate wild-type at that residue. Mutations are shaded according to frequency. **d,** Residues mutated in passaging in Huh7-ACE2 cells overlaid onto the 3CL^pro^ structure with nirmatrelvir bound. All 23 mutated residues across all the resistant populations are indicated with any individual isolate having between 1-6 mutations. The Cα of each residue that was mutated is denoted with a red sphere. The 3CL^pro^-nirmatrelvir complex was downloaded from PDB under accession code 7VH8.

Sequencing of the same wells at earlier passages revealed less diversity in 3CL^pro^, with a total of 11, 16, and 22 unique mutations detected across all populations from passages 4, 8, and 12, respectively (**Supplemental Table 1**). As a standard phylogenetic analysis showed a rather complex stepwise order of acquisition of mutations for each passaged lineage (**Extended Data Fig. 3**), we more carefully analyzed the order in which mutations arose across the various lineages (see **Methods** and **Supplemental Table 1** for details) and generated a pathway network delineating the most common routes that SARS-CoV-2 took *in vitro* to develop nirmatrelvir resistance (**Fig. 3a** **and Supplemental Table 2**). The majority of these viral lineages descended initially from T21I, P252L, and T304I, suggesting that these mutations may serve as “founder” or “precursor” mutations when the drug concentrations are relatively low. Additional mutations then occurred, probably to increase the level of resistance as the drug concentrations were increased and/or to compensate for reduced viral fitness. These findings indicated that although there are multiple solutions for SARS-CoV-2 to resist nirmatrelvir, several common mutational pathways are favored.

**Fig. 3.**
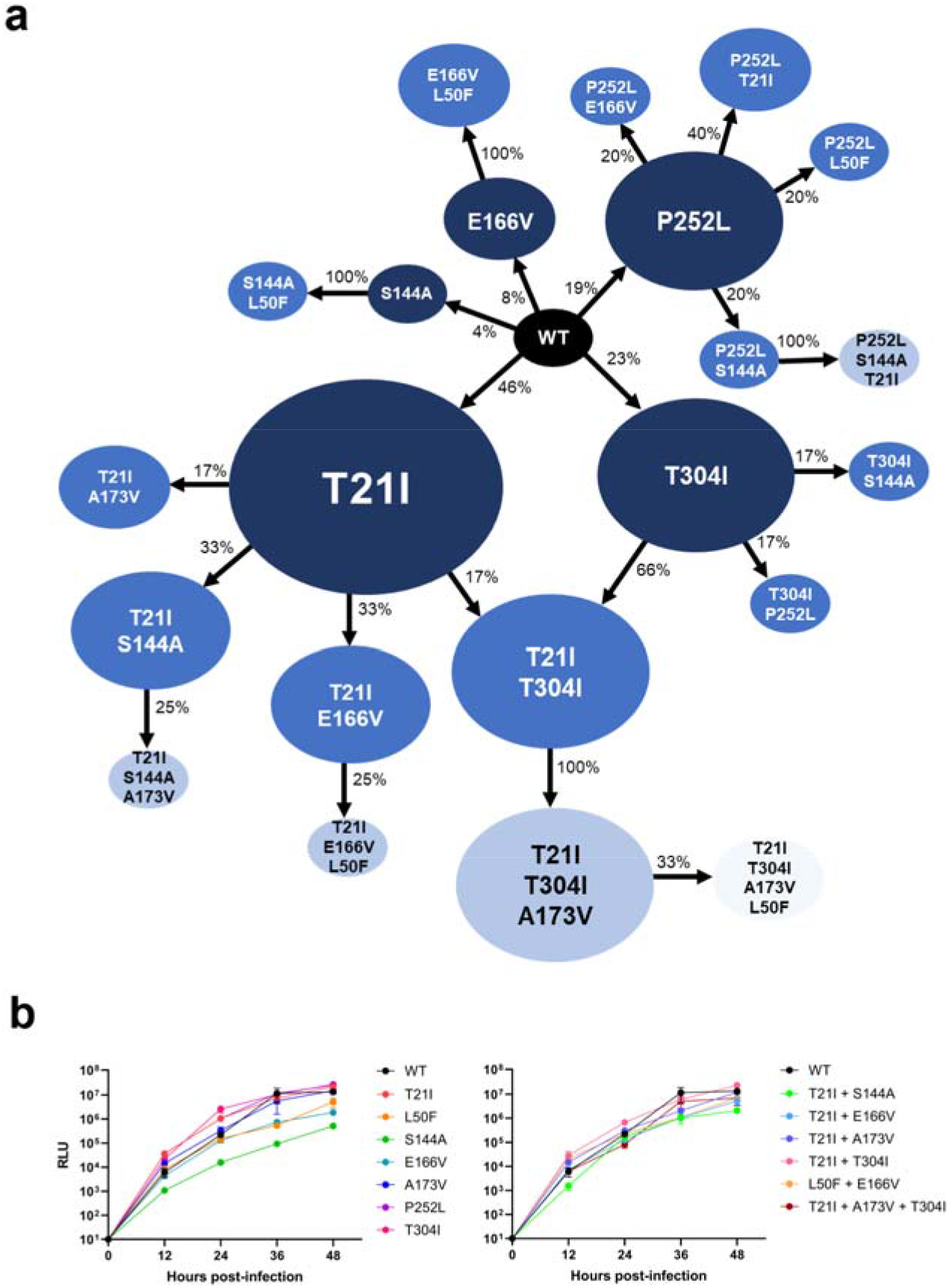
Pathways for SARS-CoV-2 resistance to nirmatrelvir. **a,** Observed pathways for nirmatrelvir resistance in Huh7-ACE2 cells. The most commonly observed mutations in passage 16 were used to build these pathways (see **Methods and Supplemental Table 2** for additional details). Nodes are colored from dark to light, with founder mutations colored darker. Percentages indicate the frequency by which the child nodes derive from the immediate parental node. Descendent arrows that do not sum to 100% indicate that a proportion did not advance beyond the indicated mutations in the experiment. **b,** Growth assay with recombinant live SARS-CoV-2 carrying single and combination 3CL^pro^ mutations. Huh7-ACE2 cells were infected with 0.01 MOI of virus and luminescence was quantified at the indicated time points. S144A, E166V, and T21I + S144A are statistically significant from WT at 48 h (two-way ANOVA with Geisser-Greenhouse correction followed by Dunnett’s multiple comparisons test; P=0.0039, P=0.0006, P=0.0006, respectively). Representative curves from a single experiment from two biologically independent experiments are shown. Error bars denote mean ± s.e.m of three technical replicates.

### Mutations conferring nirmatrelvir resistance

To further investigate which mutations were responsible for nirmatrelvir resistance, we proceeded to generate recombinant SARS-CoV-2 clones, each containing a unique mutation or a combination of mutations. To construct the 15 mutant viruses from the first passage experiment (**Fig. 1f**) and the 22 mutant viruses from the second passage experiment (**Fig. 3a**) would be beyond the scope of the current study. We therefore decided to focus on the seven most common single point mutants from the large passaging study, as well as five double mutants and one triple mutant (**Extended Data Fig. 4**). All viruses grew similarly to wild type (WT) in the absence of drug, except for S144A, E166V, and T21I + S144A, which were significantly impaired in their growth kinetics (**Fig. 3b**). However, both T21I + E166V and L50F + E166V replicated well with kinetics similar to WT, suggesting that T21I and L50F each compensated for the fitness loss of E166V. Of the individual mutants tested against nirmatrelvir, E166V was most resistant (100-fold), with P252L and T304I having low-level resistance (~6-fold), and S144A and A173V having minimal resistance (~3-fold or less) (**Figs. 4a, 4b** **and** **Extended Data Fig. 5**). Combination of either T21I or L50F with E166V resulted in a virus that was substantially resistant to nirmatrelvir (83-fold and 53-fold, respectively), but with WT replicative kinetics (**Fig. 3b**).

**Fig. 4.**
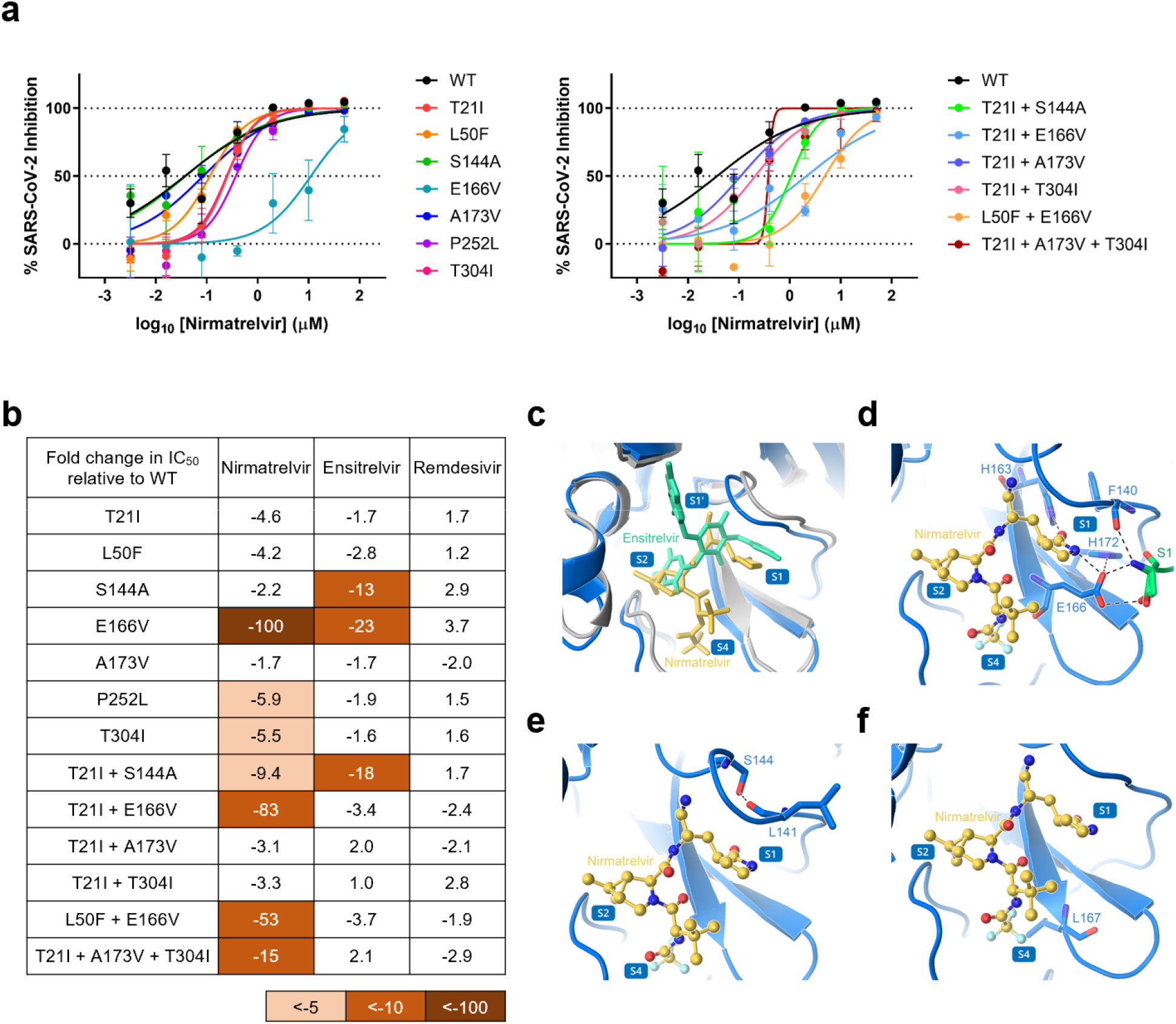
Validation of identified mutations in isogenic recombinant SARS-CoV-2. **a,** Individual inhibition curves of recombinant live SARS-CoV-2 carrying single and combination 3CL^pro^ mutations by nirmatrelvir. Representative curves from a single experiment from three biologically independent experiments are shown. Error bars denote mean ± s.e.m of three technical replicates. **b,** Inhibition of recombinant live SARS-CoV-2 carrying single and combination 3CL^pro^ mutations by nirmatrelvir, ensitrelvir, and remdesivir. Values shown are fold change of mean values in IC_50_ relative to inhibition of wild-type from three biologically independent experiments. **c,** Overlay of nirmatrelvir and ensitrelvir binding to 3CL^pro^. **d,** Several of the residues involved in direct interaction with nirmatrelvir. **e,** Several of the residues involved in formation of the S1 subsite. **f,** Interaction of L167 with nirmatrelvir. In **c-f**, nirmatrelvir is shown in yellow, enstirelvir is shown in lime green, the 3CL^pro^-nirmatrelvir complex is shown in marine, and the 3CL^pro^-ensitrelvir complex is shown in gray. Protomer A is shown in marine and protomer B is shown in green. Hydrogen bonds are indicated as black dashes. The 3CL^pro^-nirmatrelvir complex and 3CL^pro^-ensitrelvir complex were downloaded from PDB under accession codes 7VH8 and 7VU6, respectively.

We next tested this panel of viruses against ensitrelvir^28^ (also known as S-217622), another 3CL protease inhibitor that has demonstrated clinical efficacy^29^, for cross-resistance together with remdesivir as a control. Only S144A, E166V, and T21I + S144A showed substantial (13 to 23-fold) cross-resistance to ensitrelvir (**Fig. 4b** **and** **Extended Data Figs. 5**, **6**). As expected, none of these mutations conferred resistance to remdesivir. We additionally tested the passage 30 viruses resulting from the initial selection experiment in Vero E6 cells (**Fig. 1**) against these two inhibitors. Again, all three lineages were as susceptible to remdesivir as WT, and only lineage C (L50F + F140L + L167F + T304I) showed cross-resistance to ensitrelvir (~25-fold) (**Extended Data Fig. 7**). This may be due to F140L, since L50F and T304I did not demonstrate ensitrelvir resistance (**Fig. 4b**) and L167 does not contact ensitrelvir (see below). Together, these results suggest that some mutations, such as E166V, can confer a high degree of nirmatrelvir resistance alone, while others, such as T21I, P252L, and T304I, confer only low levels of nirmatrelvir resistance individually. The degree of cross-resistance to ensitrelvir was variable among the tested mutant viruses probably due to the differences in binding of these drugs to the substrate binding site of 3CL^pro^ (**Fig. 4c**). Nevertheless, it is clear that selection for nirmatrelvir resistance can yield mutations that confer cross-resistance to other inhibitors of clinical interest as well.

To begin to understand the mechanisms underlying the resistance conferred by these mutations, we considered their structural context. Nirmatrelvir and ensitrelvir both bind within the substrate binding site, but in differing modes, which may result in the differences observed in the inhibition profiles of the mutants (**Figs. 4b, 4c**). E166 directly interacts with the lactam ring of nirmatrelvir via hydrogen bonding, and the valine substitution at this position may abrogate some of these interactions to result in the strong drug resistance observed (**Fig. 4d**). E166 is also able to form hydrogen bonds with the first residue (S1) of the neighboring protomer and therefore is involved in dimerization, which is essential for protease activity as the 3CL^pro^ functions as a homodimer^30^. The disruption of the hydrogen-bonding interactions (**Fig. 4d**) may explain the reduced fitness of the E166V mutant (**Fig. 3b**). The side chain of S144 forms a hydrogen bond with the main chain of L141 to stabilize the S1 subsite of the substrate binding site, so the S144A mutation may disorder this region and hamper the binding of both nirmatrelvir and ensitrelvir (**Fig. 4e**), although it is not clear why this requires the T21I mutation in conjunction. L167 participates in the formation of the S4 subsite, and the L167F mutation may cause a steric clash with nirmatrelvir (**Fig. 4f**). However, as ensitrelvir does not extend into the S4 subsite, this mutation may not be responsible for the cross-resistance observed in lineage C (**Extended Data Fig. 7**). As F140 interacts by π-π stacking interactions with H163, which directly interacts with both nirmatrelvir and ensitrelvir, the F140L mutation may abrogate this interaction, resulting in resistance (**Figs. 4c, 4d** **and** **Extended Data Fig. 7**). For a number of these mutations, however, it is not immediately apparent how they confer drug resistance given that they are distant from the substrate binding site where the drugs bind (**Extended Data Fig. 4**).

Finally, we compared the mutations identified in this study to clinical SARS-CoV-2 sequences reported to GISAID^31^. Nearly all of the mutations we have identified were observed among the viruses circulating in the population, albeit at low frequencies (**Extended Data Fig. 8a**). Comparing the frequencies of these mutations in periods before and after the authorization of the combination of nirmatrelvir and ritonavir (PAXLOVID™) did not show an appreciable increase in the observed mutations (**Extended Data Fig. 8b**).

## Discussion

As antibody-based interventions for SARS-CoV-2 face increasing resistance by the emergence of variants of concern, antivirals with alternative modes of action have increased in importance. Nirmatrelvir, as an oral antiviral targeting 3CL^pro^, is a therapeutic that has shown high efficacy in lowering severe disease and hospitalization in infected persons who are at high risk and not vaccinated^1,2^. Indeed, it is the most commonly used antiviral drug to treat COVID-19 today^32^.

Given the adaptations that the virus has already exhibited to other modes of treatment^3–9^, it is clinically important to understand the mechanisms by which nirmatrelvir resistance can occur. The results presented herein demonstrate that *in vitro* high-level resistance to nirmatrelvir can be readily achieved by SARS-CoV-2, and that this can occur in a multitude of ways. This finding is consistent with our prior report on the extensive plasticity of the 3CL^pro^ as discovered by deep mutational scanning^33^.

In both Vero E6 cells (**Fig. 1**) and Huh7-ACE2 cells (**Fig. 2**), multiple lineages with non-overlapping mutations evolved under increasing drug pressure, consistent with what has been seen in similar small-scale studies^24,25,34,35^. Conducting selection at scale, however, revealed that there are multiple mutational pathways to nirmatrelvir resistance but with several common trajectories preferred (**Figs. 2c, 3a**). A majority of lineages descended from viruses that acquired T21I, P252L, or T304I as an initial mutation. Recombinant SARS-CoV-2 constructed to contain each of these point mutants exhibited low-level resistance (**Figs. 4a, 4b**), suggesting that each of these precursor mutations may have allowed the virus to tolerate low concentrations of nirmatrelvir but required additional mutations as the drug pressure was increased. Notably, all three of these mutations are rather distal (> 5 Å) from nirmatrelvir (**Fig. 2d**), and their mechanism for resistance is not evident without additional studies. We note, however, that T304 corresponds to the P3 site on the nsp5/6 cleavage substrate for 3CL^pro^ of both SARS-CoV and SARS-CoV-2 (**Extended Data Fig. 2**). Although the P3 site is exposed to solvent and thus not considered to confer stringent substrate specificity, it has been shown that a suitable functional group (such as the side chain of isoleucine) at the P3 site can assist in increasing inhibitor/substrate potency and selectivity for 3CL^pro^s^36–38^. Therefore, it is possible that T304I could facilitate the binding of the nsp5/6 cleavage site or promote the autocleavage process.

Analyses with isogenic mutants also revealed that several mutations are responsible for the observed nirmatrelvir resistance, with the E166V mutation conferring the most resistance (100-fold) (**Fig. 4b**), as is being reported elsewhere^33,35^. This mutation, as well, conferred a degree of cross-resistance to ensitrelvir, another clinically relevant 3CL^pro^ inhibitor^28,29^. The mechanism of resistance of E166V is explainable since it resides in the substrate binding site, and the valine substitution disrupts its hydrogen bonding to the lactam ring of nirmatrelvir (**Fig. 4d**). However, this mutation lowered the replicative fitness of the virus in vitro (**Fig. 3b**), perhaps because of a loss of interaction with the first residue of the neighboring protomer in dimerization (**Fig. 4d**)^30^. Importantly, replicative fitness was restored when T21I or L50F was added (**Fig. 3b**), without a significant impact on drug resistance (**Fig. 4b**). How these two mutations compensate for the fitness loss of E166V remains unknown. It is worth mentioning that the E166V mutation was reported to be found in viral isolates from a few of the PAXLOVID™-treated individuals in the EPIC-HR clinical trial^1^ (see Fact Sheet for Healthcare Providers: Emergency Use Authorization for PAXLOVID™, revised July 6, 2022).

We have also found that a number of additional mutations could confer resistance to nirmatrelvir *in vitro*. T21I + S144A mediated not only significant resistance to nirmatrelvir but also cross-resistance to ensitrelvir (**Fig. 4b**), but this virus exhibited slower growth kinetics (**Fig. 3b**). Likewise, we inferred that both L167F and F140L were likely mediating drug resistance in the C-P30 lineage of the first *in vitro* passaging experiment (**Fig. 1f**) as discussed above along with possible structural explanations. It is clear, nevertheless, that we have only studied a limited number of the mutational pathways that SARS-CoV-2 took to evade nirmatrelvir. Furthermore, many of the mutations revealed by our study are without a straightforward structural explanation at this time. It should also be mentioned that our studies were conducted with the ancestral WA1 strain, and the currently circulating Omicron variants, all of which except for BA.3 contain a P132H mutation in 3CL^pro^, may differ in their nirmatrelvir evasion pathways. While this mutation has been reported to have no direct effect on nirmatrelvir resistance, it may influence the emergence of subsequent resistance conferring mutations^39^. It will require extensive virological, biochemical, and structural studies to delineate which mutations confer resistance and how, as well as to understand how certain mutations play compensatory roles. A better understanding of the mechanisms of nirmatrelvir resistance could provide insight into the development of the next generation of 3CL^pro^ inhibitors.

Nirmatrelvir has been used to treat COVID-19 for only 6 months or less in most countries. SARS-CoV-2 resistance to this drug in patients has yet to be reported, and we see no appreciable difference in frequencies of the 3CL^pro^ mutations that we have uncovered in periods before and after the emergency use authorization (**Extended Data Fig. 8**). Perhaps the lack of nirmatrelvir resistance in patients to date is due to the high drug concentrations achieved with the prescribed regimen, making it difficult for the virus to accumulate mutations in a stepwise manner. In addition, the drug is administered while the immune system is also actively eliminating the virus, including any resistant forms that may have emerged. Therefore, it makes sense to focus our surveillance effort on immunocompromised individuals on nirmatrelvir treatment for the appearance of drug-resistant virus. Past experience with other viral infections tells us that if drug resistance could be selected *in vitro*, it surely will occur *in vivo*. Although current COVID-19 therapies have been largely administered as monotherapies, it is possible that future treatment will benefit from the use of a combination of drugs to minimize the likelihood of SARS-CoV-2 escape.

## Figure Legends

**Extended Data Fig. 1.**
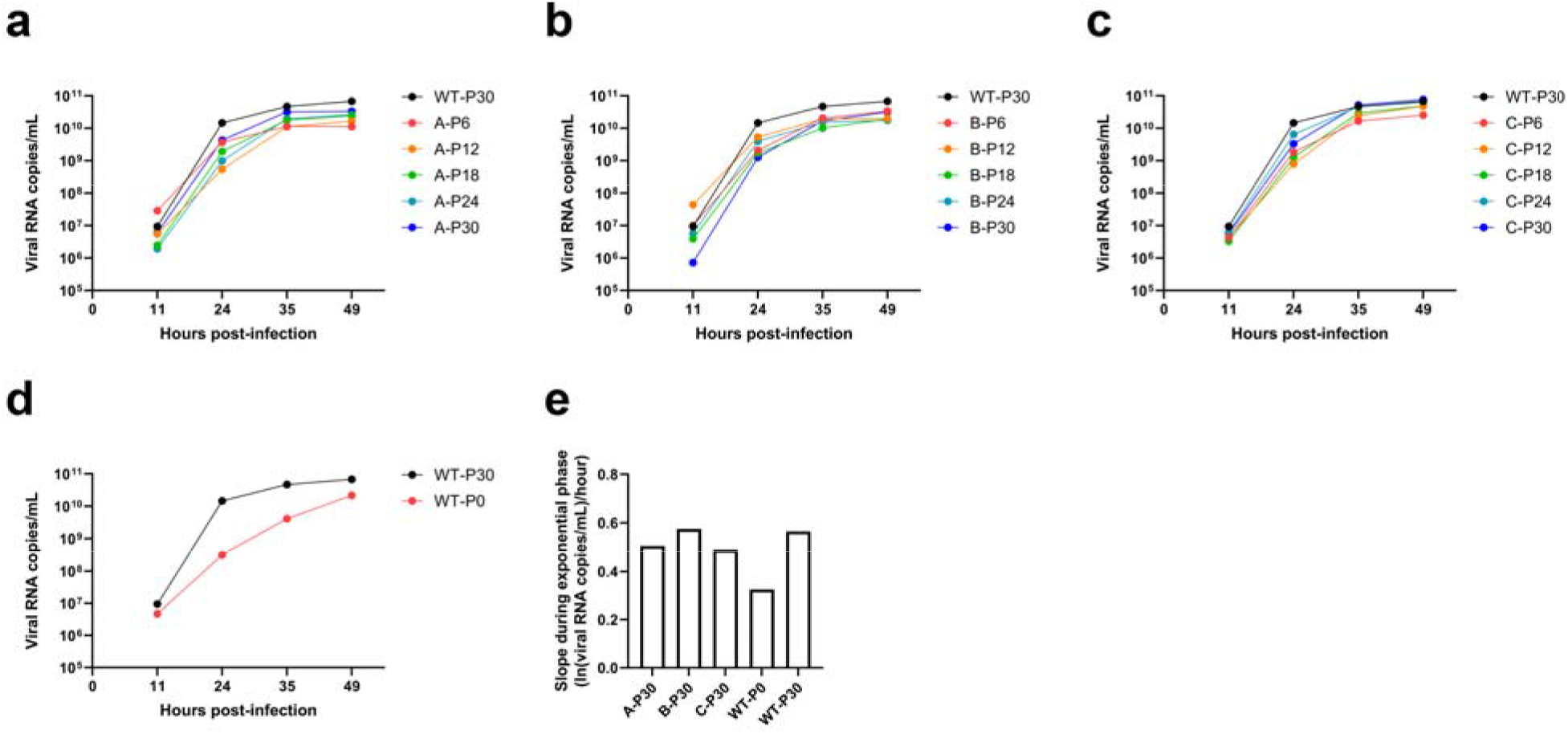
Growth assays with SARS-CoV-2 passaged in Vero E6 cells. **a-d,** Growth was quantified for lineage A (**a**), lineage B (**b**), lineage C (**c**), and unpassaged SARS-CoV-2 (**d**, denoted as WT-P0) in comparison to SARS-CoV-2 passaged without nirmatrelvir for 30 passages (denoted as WT-P30). Vero E6 cells were infected with 200 TCID_50_ of the indicated viruses and viral RNA was quantified at the indicated time points. **e,** The slope during the exponential phase (between 11 and 24 hours post-infection) of growth for the indicated viruses.

**Extended Data Fig. 2.**
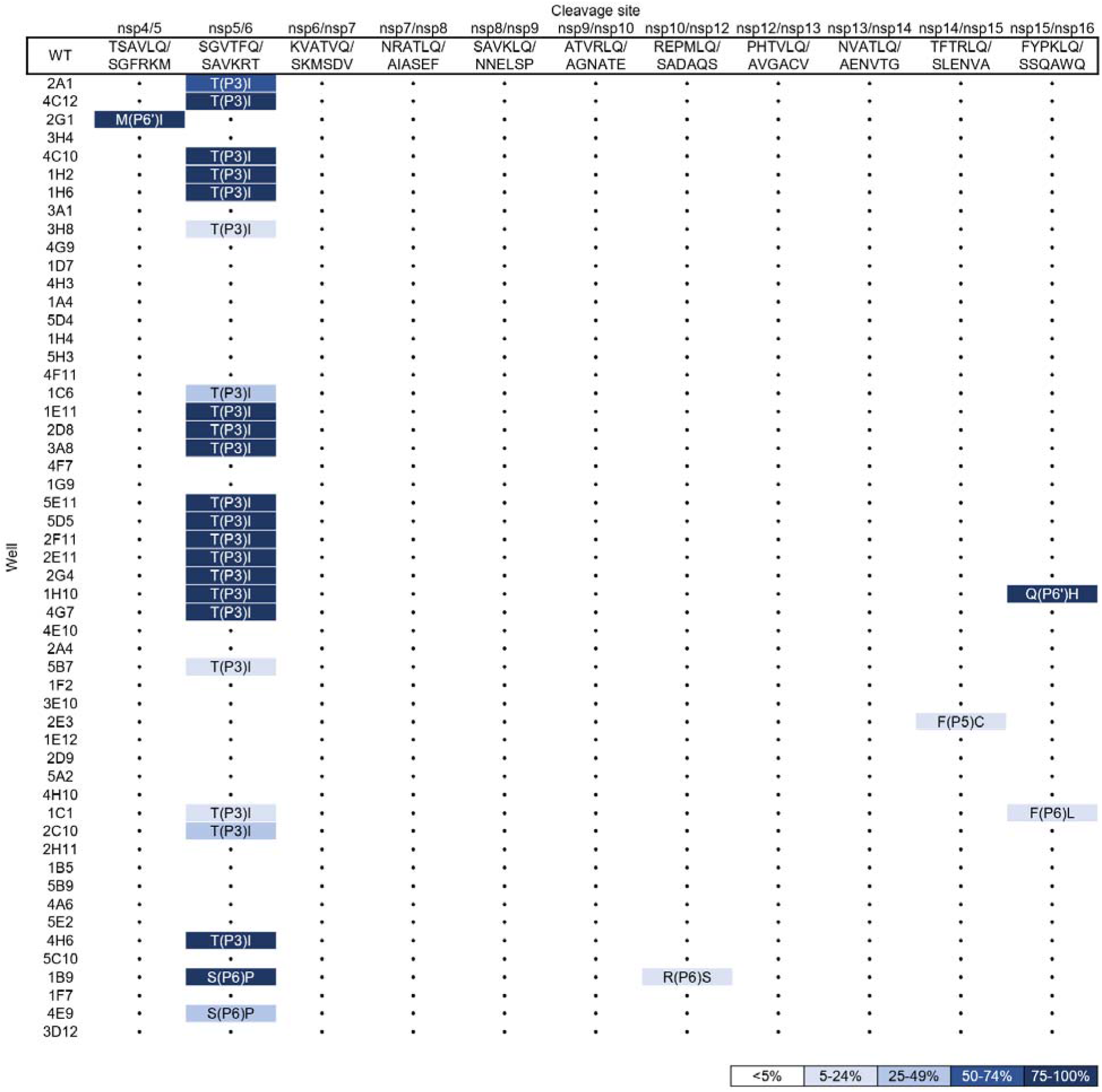
Mutations in the 11 3CL^pro^ cut sites found in passage 16 from the 53 wells passaged in Huh7-ACE2 cells. Dots indicate wild-type at that cut site.

**Extended Data Fig. 3.**
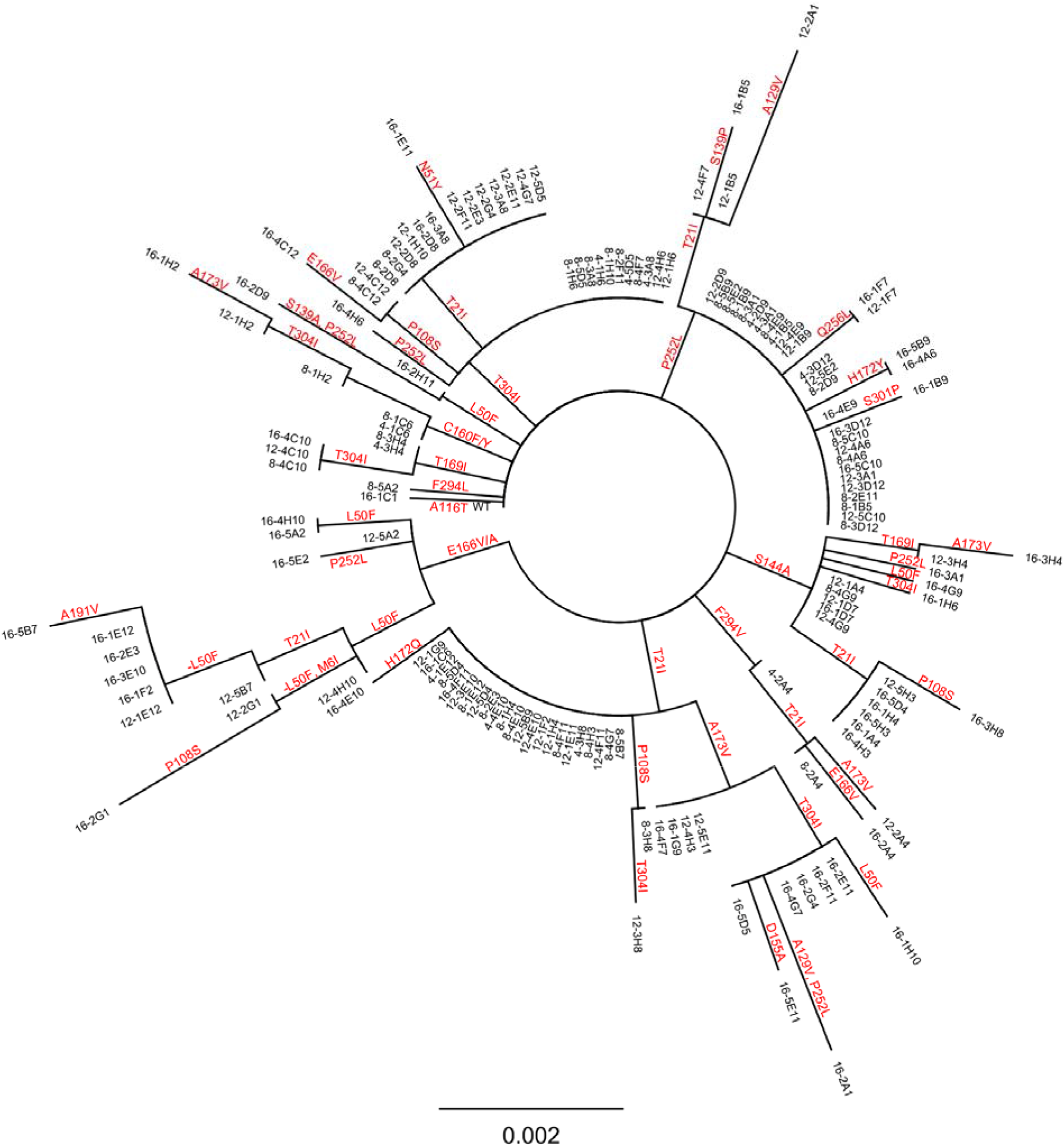
Phylogenetic tree of sequences from passaging in Huh7-ACE2 cells. Only sequences with mutations are shown. Sequences are denoted as passage number, followed by the well number. Mutations that arose along particular branches are annotated in red, “-” denotes when a mutation appears to be lost from a particular branch.

**Extended Data Fig. 4.**
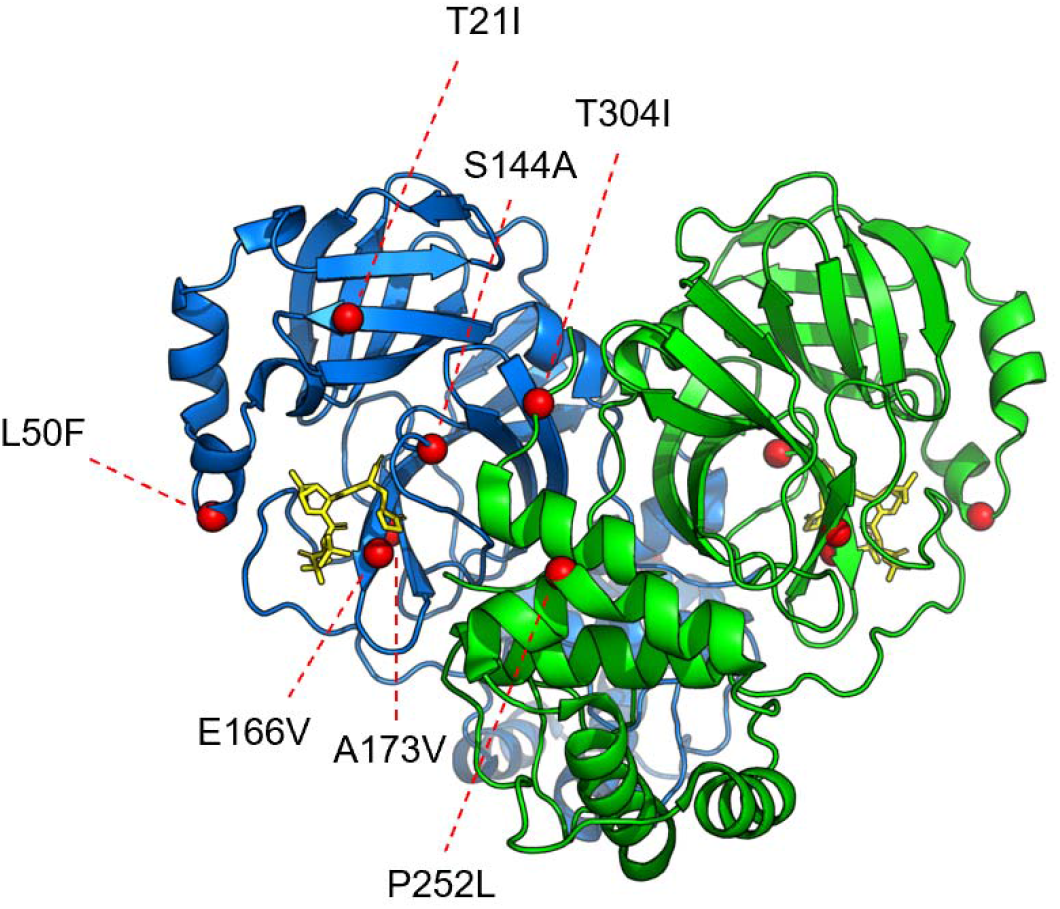
Mutations studied as isogenic recombinant SARS-CoV-2 overlaid onto the 3CL protease structure. The Cα of each residue that was mutated is denoted with a red sphere. The 3CL^pro^-nirmatrelvir complex was downloaded from PDB under accession code 7VH8.

**Extended Data Fig. 5.**
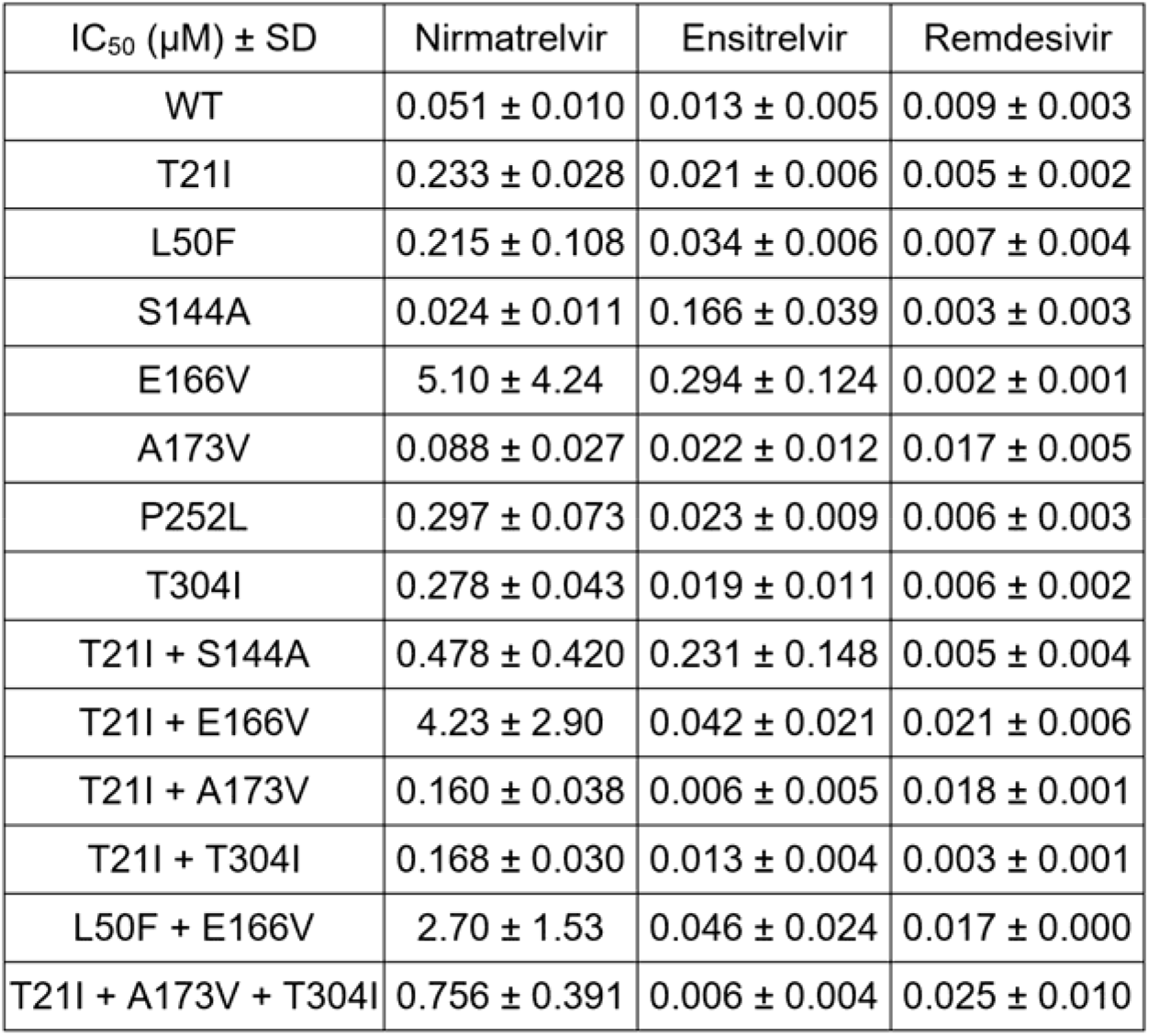
Raw IC_50_ values for recombinant live SARS-CoV-2 carrying single and combination 3CL^pro^ mutations by nirmatrelvir, ensitrelvir, and remdesivir. Mean ± SD of three biologically independent experiments are shown.

**Extended Data Fig. 6.**
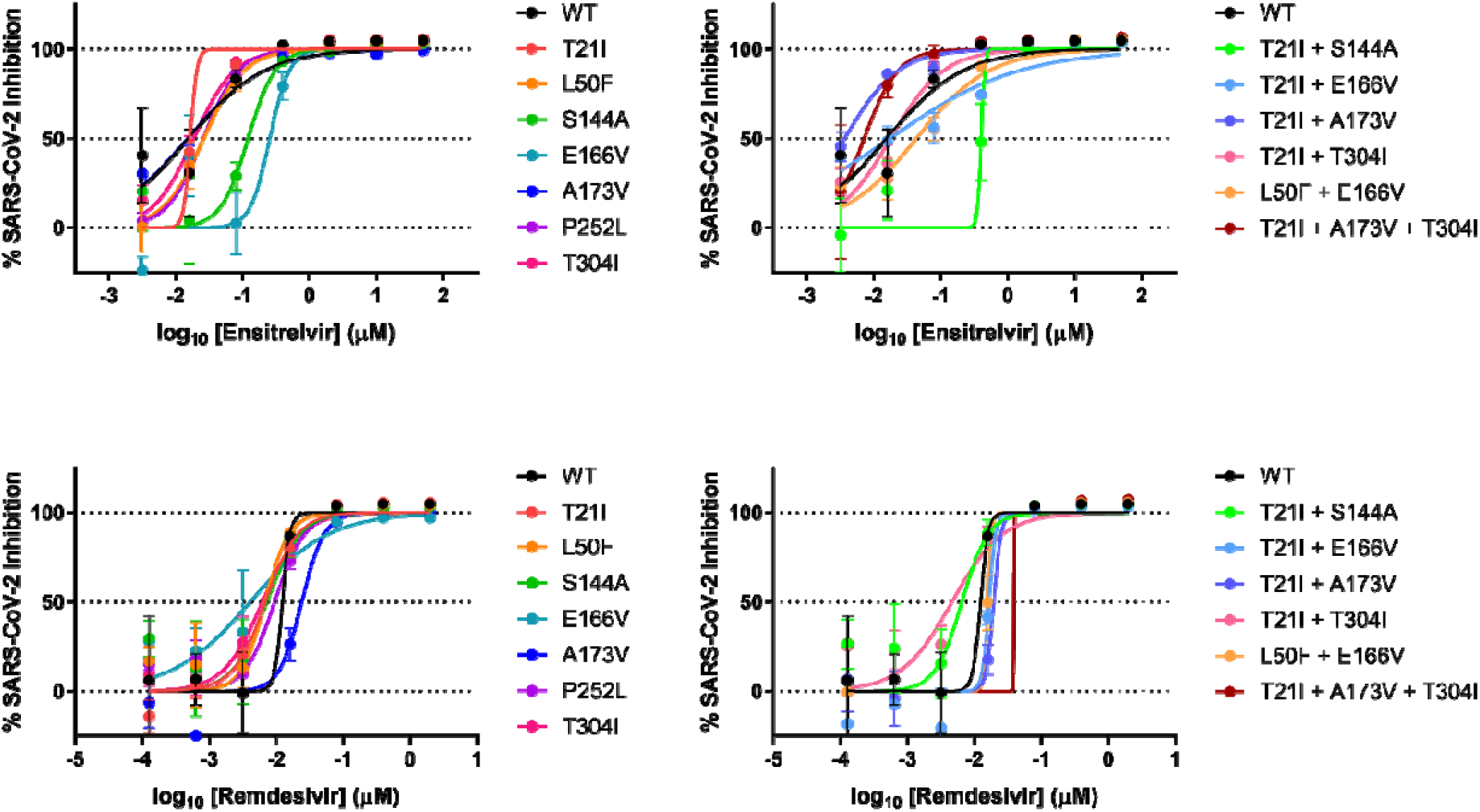
Individual inhibition curves of recombinant live SARS-CoV-2 carrying single and combination 3CL^pro^ mutations by ensitrelvir and remdesivir. Representative curves from a single experiment from three biologically independent experiments are shown. Error bars denote mean ± s.e.m of three technical replicates.

**Extended Data Fig. 7.**
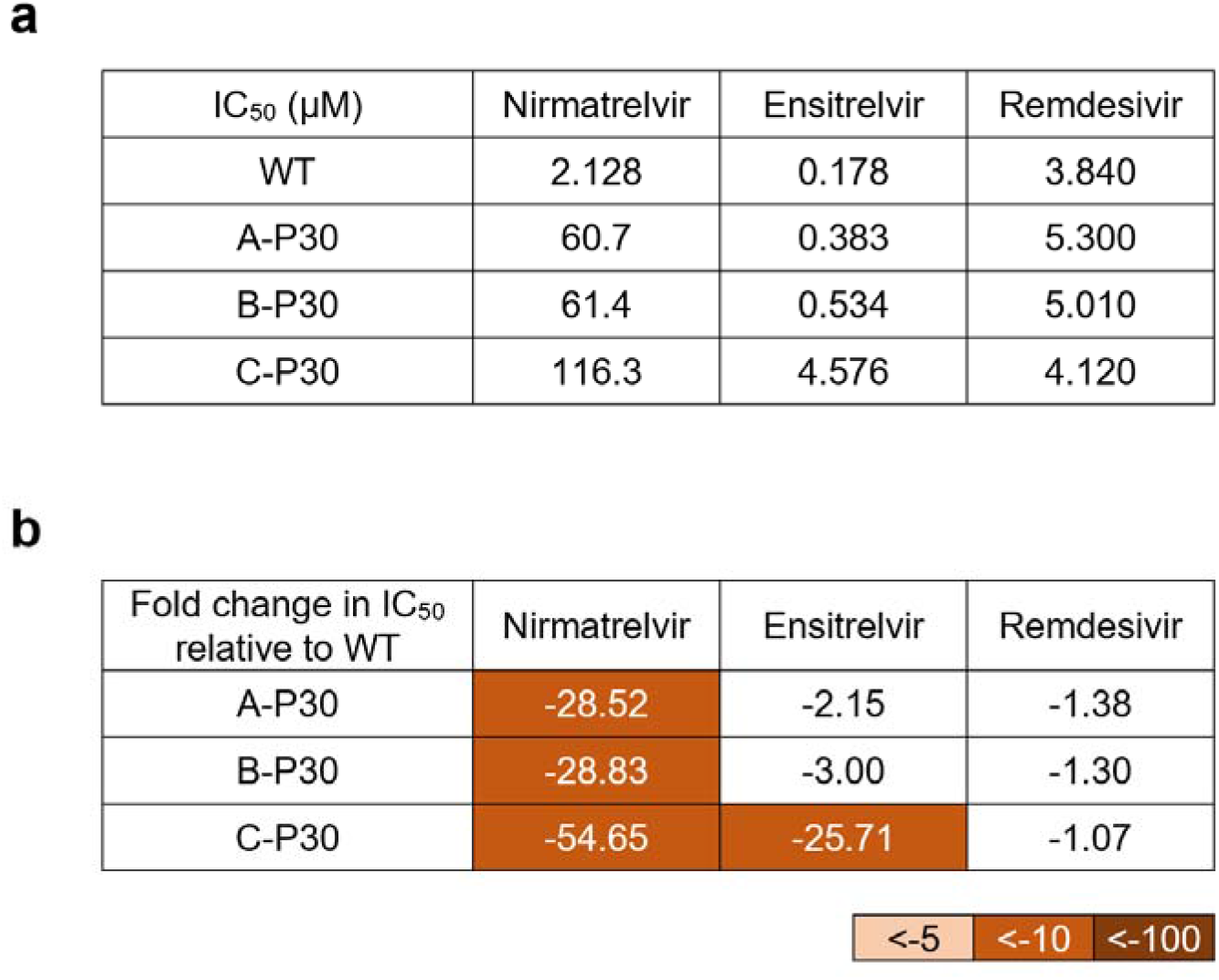
Inhibition of passage 30 of SARS-CoV-2 passaged in Vero E6 cells by nirmatrelvir, ensitrelvir, and remdesivir. **a,** Raw IC_50_ values. **b,** Fold change relative to inhibition of wild-type.

**Extended Data Fig. 8.**
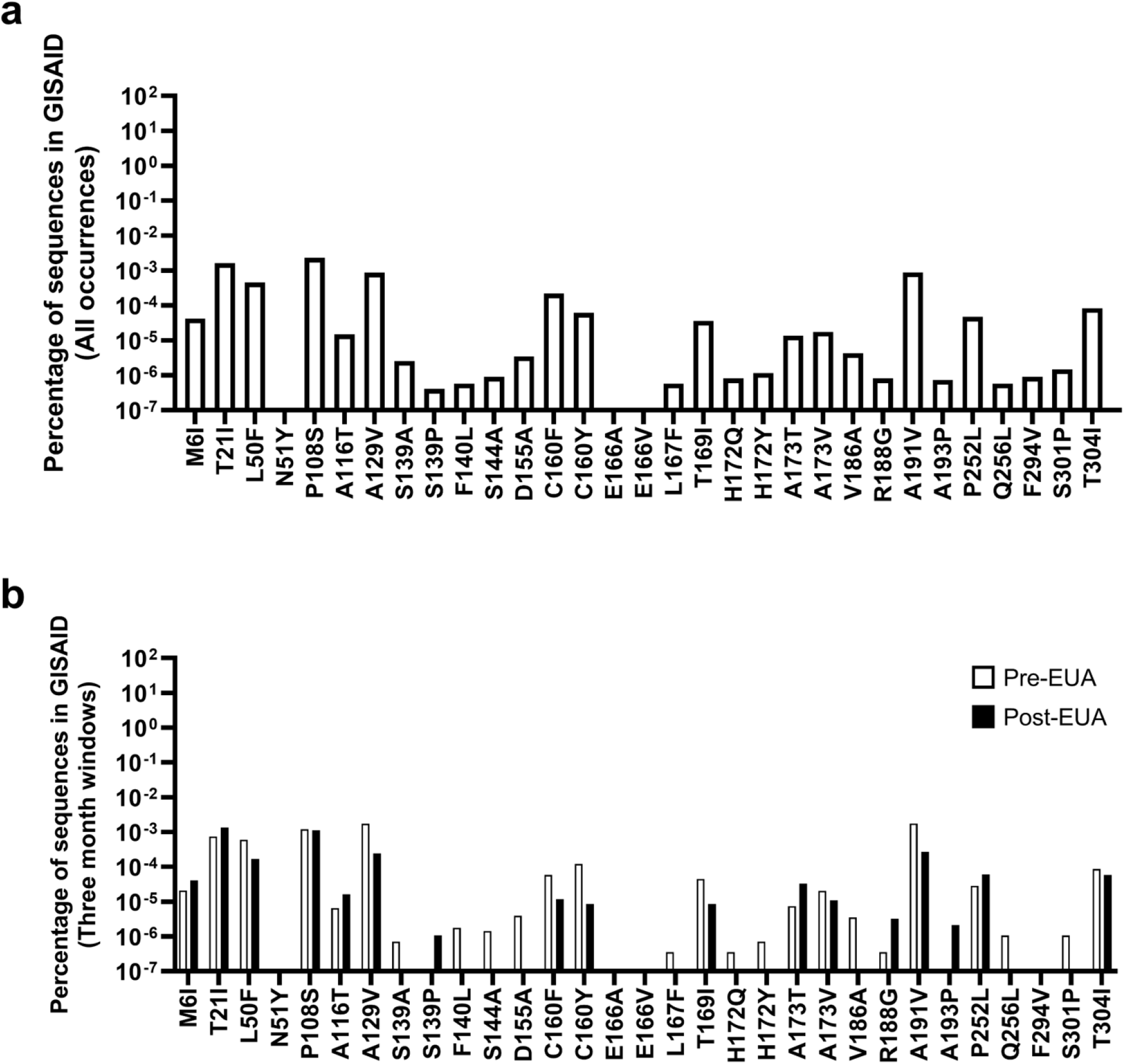
Frequencies of identified 3CL^pro^ mutations in GISAID. **a,** All occurrences of the indicated mutations were tabulated from GISAID. **b,** All occurrences of the indicated mutations were tabulated from GISAID in the three months prior to EUA (9/22/2021 to 12/22/2021) or after EUA (3/26/2022 to 6/26/2022).

**Supplemental Table 1. Raw sequencing results for passaging in Huh7-ACE2 cells.** Samples are denoted as passage number, followed by well number. Cut site mutations are denoted as “cs”.

**Supplemental Table 2. Passage transitions used for construction of the pathway analysis in Figure 3a.**

**Supplemental Table 3. GenBank Accession IDs for sequences from passaging in Vero E6 cells.**

**Supplemental Table 4. Oligos used for next-generation sequencing.**

**Supplemental Table 5. SRA Accession IDs for raw sequencing data from passaging in Huh7-ACE2 cells.**

**Supplemental Table 6. Oligos used for site-directed mutagenesis to produce isogenic recombinant SARS-CoV-2.**

## Methods

### Biosafety

All SARS-CoV-2 passaging, infection, and recombinant virus production was conducted in BSL-3 laboratories at Columbia University Irving Medical Center under procedures and guidelines approved by the Columbia University Institutional Biosafety Committee (IBC).

### Compounds

Nirmatrelvir was purchased from Aobius, ensitrelvir was purchased from Glixx Laboratories, and remdesivir was purchased from Selleckchem.

### Cells

Vero E6 cells were obtained from ATCC (Catalog #CRL-1586), HEK293T cells were obtained from ATCC (Catalog #CRL-3216), and Vero E6-TMPRSS2-T2A-ACE2 cells were obtained from BEI Resources (Catalog #NR-54970). Huh7-ACE2 cells were generated previously^33,40^. Cell morphology was visually confirmed prior to use and all cell lines tested mycoplasma negative. All cells were maintained at 37 °C under 5% CO_2_.

### In vitro selection for SARS-CoV-2 resistance to nirmatrelvir in Vero E6 cells

To select for the development of drug resistance against nirmatrelvir, WA1 (SARS-CoV-2, USA-WA1/2020 strain) was cultured in the presence of increasing concentrations of nirmatrelvir and passaged 30 times. Virus isolates recovered from the culture at various passages were then characterized for their resistance to nirmatrelvir and their replication capacity.

To initiate the passaging, Vero E6 cells were seeded in a 24 well-plate at a density of 1 × 10^5^ cells per well in complete media (DMEM + 10% FCS + penicillin/streptomycin), and then the drug and virus were added the following day. The drug was prepared in a three-fold dilution series based on the original IC_50_ of the drug. The virus was added at 5,000 TCID_50_ per well. Three days post-infection, each well was scored for cytopathic effects (CPE) in a range of 0 to 4+ based on comparison to control wells as previously described^41^, and 100 μL of the supernatant from the well with a CPE score equal to or greater than 2+ was passaged to each well in the next culture plate. The passage culture was set up in triplicate (lineages A, B, and C) and the passaging was performed independently, i.e., viruses in lineage A were kept within the lineage A series of wells at every passage. Along with the cultures passaged with nirmatrelvir, WA1 was passaged without nirmatrelvir in two independent wells to serve as a passage control.

IC_50_s for each lineage in the passaging were determined based on the CPE scores at day 3 of each passage. IC_50_ values were derived by using DeltaGraph (Red Rock Software).

### Sequencing of SARS-CoV-2 passaged in Vero E6 cells

For the SARS-CoV-2 passaged in Vero E6 cells, passages were sequenced by Sanger sequencing or by Nanopore sequencing. For Sanger sequencing, viral RNA was isolated from the culture supernatant with QIAamp® Viral RNA Mini Kit (Qiagen), reverse transcribed to cDNA with Superscript IV™ Reverse Transcriptase (Thermo Fisher) and the priming primer, nsp5.R1, and subjected to nested PCR with Platinum™ SuperFi II (Thermo Fisher) to obtain the full length nsp5 gene. The primers for the first PCR are nsp5.F1: 5’-GTAGTGATGTGCTATTACCTCTTACGC-3’ and nsp5.R1: 5’-GCAAAAGCAGACATAGCAATAATACC-3’. The primers for the second PCR are nsp5.F2: 5’-CTTCAGTAACTCAGGTTCTGATGTTCT-3’ and nsp5.R2: 5’-ACCATTGAGTACTCTGGACTAAAACTAAA-3’. Both PCRs were run with the same condition of 98 °C for 30 s, 25 cycles of 98 °C for 15 s, 60 °C for 10 s, and 72 °C for 1 min, followed by 72 °C for 5 min. The PCR products were purified and sequenced (Genewiz). Mixtures of viruses were determined by inspecting sequencing chromatograms. The sequences were analyzed using Lasergene software (DNASTAR).

For Nanopore sequencing, viral RNA was isolated from the culture supernatant with QIAamp® Viral RNA Mini Kit (Qiagen), and then the Midnight RT PCR Expansion kit and Rapid Barcoding kits (Oxford Nanopore) were used to amplify and barcode overlapping 1,200 bp amplicons tiled across the viral genome^42,43^. An Oxford Nanopore GridION with R9.4.1 flow cells was used for sequencing. Basecalling was performed in MinKNOW v22.05.1. Consensus sequence generation was performed using the ONT Epi2Me ARTIC Nextflow pipeline v0.3.16 (https://github.com/epi2me-labs/wf-artic). Pangolin 4.0.6 with UShER v1.6 was used for parsimony-based lineage assignment. Sequences have been deposited to GenBank (ON924329-ON924335, ON930401-ON930431) (**Supplemental Table 3**).

### Inhibition assay with SARS-CoV-2 passaged in Vero E6 cells

To characterize the inhibition of passaged viruses, each of the viruses were first propagated in Vero E6 cells in the absence of drug and titrated by the Reed-Muench method^44^. Vero E6 cells were then seeded in 96 well-plates at a density of 1.5 × 10^4^ cells per well in complete media. The following day, the virus was inoculated at a dose of 500 TCID_50_ per well, and a two-fold dilution series of inhibitor was added in quadruplicate. Three days post-infection, the level of CPE was scored and the IC_50_ was derived by fitting a nonlinear regression curve to the data in GraphPad Prism version 9.4 (Dotmatics).

### Growth assay with SARS-CoV-2 passaged in Vero E6 cells

The fitness of passaged viruses was characterized by a viral growth assay. Vero E6 cells were seeded in 96 well-plates at a density of 1.5 × 10^4^ cells per well in complete media. The following day, the virus was inoculated at a dose of 200 TCID_50_ per well in quadruplicate. At 6 h post-infection, free virions in the culture were removed by changing of the media twice. At 11, 24, 35, and 49 h post-infection, 50 μL of the culture supernatant from each well was collected and replenished with an equivalent volume of fresh media. Viral RNA from each time point was purified by using PureLink™ Pro 96 Viral RNA/DNA Purification Kit (Thermo Fisher) and then the viral copy number in each sample was estimated by qRT-PCR using TaqPath™ 1-Step RT-qPCR Master Mix (Thermo Fisher) and 2019-nCov CDC EUA Kit (Integrated DNA Technologies) with 7500 Fast Dx Real-Time PCR Instrument (Applied Biosystems).

### In vitro selection for SARS-CoV-2 resistance to nirmatrelvir in Huh7-ACE2 cells

To conduct selection at scale to observe as many resistance pathways as possible, SARS-CoV-2 infection was conducted in five 96 well-plates, thereby allowing for 480 independent selection lineages. We hypothesized that the use of limited number of cells allowed for a “bottleneck effect” to occur, which enabled observation of rarer events that may be outcompeted from a larger population.

To initiate the passaging, 3 × 10^4^ Huh7-ACE2 cells per well were seeded in complete media in five 96 well-plates. The following day, all wells were infected with 0.05 MOI of SARS-CoV-2-mNeonGreen (a fluorescent reporter variant of USA-WA1/2020, gift of Pei-Yong Shi)^27^ without drug to generate passage 0 (P0). For each successive passage, cells were seeded the day prior to infection, and then the drug and virus were added, three to four days post-infection of the previous passage. The drug was initially added at 25 nM and then doubled every other successive passage. Viruses were transferred between passages by overlaying 50 μL of the supernatant from the previous passage. After 16 passages, all 54 wells positive for mNeonGreen signal were sequenced, of which 53 lineages could be determined.

### Inhibition assay with SARS-CoV-2 passaged in Huh7-ACE2 cells

To characterize the inhibition of passaged viruses, each of the viruses were first propagated in Huh7-ACE2 cells in the absence of drug and titrated by the Reed-Muench method^44^. Huh7-ACE2 cells were then seeded in 96 well-plates at a density of 2 × 10^4^ cells per well in complete media. The following day, the virus was inoculated at a dose of 0.05 MOI per well, and a five-fold dilution series of inhibitor was added in triplicate. At 24 h post-infection, the supernatant was aspirated and cells were fixed with 4% PFA in PBS and stained with DAPI. Cells were then imaged for DAPI and GFP using IN Cell 2000 (GE) and analyzed with CellProfiler version 4.0.7 ^45^. The IC_50_ was then derived by fitting a nonlinear regression curve to the data in GraphPad Prism version 9.4 (Dotmatics).

### Sequencing of SARS-CoV-2 passaged in Huh7-ACE2 cells

For the SARS-CoV-2 passaged in Huh7-ACE2 cells, passages were sequenced by Illumina next-generation sequencing. Viral RNA was first extracted using PureLink™ Pro 96 Viral RNA/DNA Purification Kit (Thermo Fisher). Reverse transcription was carried out using Maxima H Minus First Strand cDNA Synthesis Kit (Thermo Fisher) with random hexamers according to the manufacturer’s instructions. Briefly, 13.75 μL of viral RNA was mixed with 0.25 μL random hexamers (50 ng/μL) and 1 μL dNTPs (10 mM), and incubated at 65 °C for 5 min followed by 4 °C for 1 min. Then, a mixture containing 4 μL 5x RT buffer, 0.25 μL enzyme mix (containing Maxima H Minus RT and RNase inhibitor), and 0.75 μL H_2_O was added to each sample and the reactions were incubated at 25 °C for 10 min, 55 °C for 30 min, and 85 °C for 5 min.

Sequencing libraries were prepared by amplifying either nine fragments tiled across the 3CL^pro^ open reading frame and adjacent nsp4/5 and nsp5/6 cut sites, or nine fragments containing each of the remaining 3CL^pro^ cut sites (see **Supplemental Table 4** for primer sequences). Primers amplifying non-adjacent fragments of the 3CL^pro^ were pooled together and reactions were carried out in technical duplicate, for a total of four first-round PCRs per sample. Each first-round PCR contained the following components: 1 μL cDNA, 0.25 μL 100 μM pooled primers, 0.4 μL 10 mM dNTPs, 2 μL 10x Taq buffer, 0.1 μL Taq DNA polymerase (Enzymatics), and 16.25 μL H_2_O. Cycling conditions were as follows: (1) 94 °C, 3 min, (2) 94 °C, 30 s, (3) 54 °C, 20 s, (4) 72 °C, 30 s, (5) Return to step #2 for 34 additional cycles, (6) 72 °C, 3 min, (7) Hold at 4 °C.

Products from the four first-round PCRs for each sample were pooled and gel purified, and a second-round indexing PCR was carried out for each sample with the following reagents: 1 μL template DNA, 0.25 μL each 100 μM indexing primer, 0.4 μL 10 mM dNTPs, 2 μL 10x Taq buffer, 0.1 μL Taq DNA polymerase, and 16.25 μL H_2_O. The cycling conditions were as follows: (1) 94 °C, 3 min, (2) 94 °C, 30 s, (3) 54 °C, 20 s, (4) 72 °C, 30 s, (5) Return to step #2 for 6 additional cycles, (6) 72 °C, 3 min, (7) Hold at 4 °C.

Second round PCR products were pooled, gel purified, and sequenced on an Illumina NextSeq system with 150 bp single-end reads. For select samples, sequences were confirmed using nanopore sequencing (Plasmidsaurus). For samples P16-2D9, P12-1A4, and 4-3A1, the original Illumina sequencing results were replaced with the Nanopore sequencing results.

For each sample, mutations and their frequencies were identified using the V-pipe computational pipeline (version 2.99.2)^46^, with Wuhan-Hu-1 (GenBank accession no. MN908947) set as the reference sequence. Frequency thresholds for reporting mutations were set at 5% and 10% for Illumina and nanopore sequencing, respectively. See **Supplemental Table 1** for absolute frequencies of mutations within each sample. Raw sequencing data have been deposited to NCBI Short Read Archive under BioProject Accession ID PRJNA852265 (see **Supplemental Table 5** for SRA Accession IDs for each sample). These sequences were clustered for **Fig. 2c** using seaborn.clustermap under default settings, which utilizes the UPGMA algorithm through SciPy^47,48^. The phylogenetic analysis shown in **Extended Data Fig. 3** was produced in Geneious Prime v2022.1 (Dotmatics) with PHYML extension, using the GTR substitution model with the optimization conditions of topology/length/rate.

### Pathway analysis for SARS-CoV-2 passaged in Huh7-ACE2 cells

Fig. 3a was constructed from lineages containing only the mutations that were found most commonly in passage 16: T21I, T304I, A173V, E166V, P252L, S144A, and L50F. These lineages were determined based on the frequencies of the corresponding mutations in a given well at each passage. Pairs of mutants whose frequencies summed to greater than 100% were assumed to co-occur on the same allele. The same logic was extended to identify triple and quadruple mutants, such that if each pairwise sum of frequencies within a group of mutations was greater than 100%, all mutations within that group were assumed to occur together. The order in which mutations in a given lineage arose was imputed either from stepwise appearance over time (e.g., passage 4 has mutation 1 and passage 8 has mutation 1 and mutation 2 at a total combined frequency >100%) with increasing frequencies, or, in cases where 2 mutations arose between sequenced passages and were deemed to co-occur in a single virus, by their relative frequencies (e.g., if passage 4 has no mutations and passage 8 has mutation 1 at 99% frequency and mutation 2 at 30% frequency, mutation 1 was assumed to have arisen first). See **Supplemental Table 2** for the datapoints used in this analysis.

### Recombinant SARS-CoV-2 production

A reverse genetics system based on the pBeloBAC11 bacterial artificial chromosome (BAC) containing the SARS-CoV-2 genome with a NanoLuc luciferase reporter replacing ORF7a^49^ (gift of Luis-Martinez Sobrido) was used to produce recombinant SARS-CoV-2 harboring 3CL^pro^ mutations. Mutants BACs were produced as previously described^33^; see **Supplemental Table 6** for a list of mutagenic primers used. These BACs (2 μg each) were then transfected into HEK293T cells in 12 well-plates in triplicate using Lipofectamine™ 3000 Transfection Reagent (Thermo Fisher) according to the manufacturer’s instructions. Two days post-transfection, cells were pooled and overlaid onto Vero E6-TMPRSS2-T2A-ACE2 cells in 25 cm^2^ flasks. After three days, the supernatant was collected from these cells and clarified by centrifugation, then used to infect Vero E6 cells in 75 cm^2^ flasks. Four days post-infection, the supernatant was harvested, clarified by centrifugation, and aliquoted. Viruses were stored at −80 °C prior to use. An aliquot of all recombinant viruses was confirmed by nanopore sequencing for the mutation of interest and for purity prior to use.

### Inhibition assay with recombinant SARS-CoV-2

Viruses were first titrated to normalize input. To characterize inhibition, Huh7-ACE2 cells were seeded at a density of 2 × 104 cells per well in 96 well-plates. The following day, cells were infected with 0.05 MOI of virus, and treated with inhibitor in a five-fold dilution series. One day post-infection, cells were lysed and luminescence was quantified using the Nano-Glo® Luciferase Assay System (Promega) according to the manufacturer’s instructions. IC50s were derived by fitting a nonlinear regression curve to the data in GraphPad Prism version 9.4 (Dotmatics).

### Growth assay with recombinant SARS-CoV-2

Viruses were first titrated to normalize input. To characterize fitness, Huh7-ACE2 cells were seeded at a density of 2 × 10^4^ cells per well in 96 well-plates. The following day, cells were infected with 0.01 MOI of virus. At 12, 24, 36, and 48 h post-infection, cells were lysed and luminescence was quantified using the Nano-Glo® Luciferase Assay System (Promega) according to the manufacturer’s instructions.

### Retrieval of clinical mutation frequencies

COVID-19 CG was used to retrieve all clinically observed 3CL^pro^ mutations from GISAID on June 26, 2022, either since the start of the COVID-19 pandemic or between March 26-June 26, 2022, and September 22-December 22, 2021^31,50^.

## Acknowledgements

This study was supported by funding from the JPB Foundation, Andrew and Peggy Cherng, Samuel Yin, Carol Ludwig, and David and Roger Wu to D.D.H. A.C. is supported by a Career Awards for Medical Scientists from the Burroughs Wellcome Fund. We thank Pei-Yong Shi for the SARS-CoV-2-mNeonGreen reporter virus and Luis Martinez-Sobrido and Chengjin Ye for the bacterial artificial chromosome system to generate recombinant SARS-CoV-2.

## Author contributions

S.I., B.C., S.J.H., A.C., and D.D.H. conceived this project and approach to scaled screening for resistant viral variants. S.I. and H.M. conducted the *in vitro* passaging. S.I., H.M., B.C., S.J.H., M.K.A., A-C.U., and A.C. conducted the sequencing. S.I. and H.M. conducted the fitness and inhibition assays. B.C., S.J.H., and A.C. conducted the pathway analyses. S.I., H.M., B.C., S.J.H., M.I.L., Y.S., and A.C. generated recombinant SARS-CoV-2. Y.D., Y.G., Z.S., and H.Y. conducted the structural analyses. S.P.G. contributed to discussions of the data and analysis. S.I., H.M., B.C., S.J.H., Y.D., H.Y., A.C., and D.D.H. wrote the manuscript with input from all authors.

## Materials availability

Materials used in this study will be made available under an appropriate Materials Transfer Agreement.

## Data availability

All experimental data are provided in the manuscript. The sequences of mutants from passaging in Vero E6 cells have been deposited to GenBank (ON924329-ON924335, ON930401-ON930431). The raw next-generation sequencing data of passaging in Huh7-ACE2 cells are available from the NCBI Short Read Archive under BioProject Accession ID PRJNA852265. The structures of the 3CL^pro^-nirmatrelvir complex and 3CL^pro^-ensitrelvir complex were downloaded from PDB under accession codes 7VH8 and 7VU6, respectively.

## Code availability

Sequencing data processing and visualization was performed using the ONT ARTIC Nextflow pipeline (https://github.com/epi2me-labs/wf-artic) and the V-pipe computational pipeline (https://github.com/cbg-ethz/V-pipe), and clustering was performed using seaborn (https://github.com/mwaskom/seaborn), which utilizes SciPy (https://github.com/scipy/scipy), all of which are publicly available software and packages.

## References

1 Hammond, J. et al. Oral Nirmatrelvir for High-Risk, Nonhospitalized Adults with Covid-19. N Engl J Med 386, 1397–1408, doi:10.1056/NEJMoa2118542 (2022).

2 Owen, D. R. et al. An oral SARS-CoV-2 M(pro) inhibitor clinical candidate for the treatment of COVID-19. Science 374, 1586–1593, doi:10.1126/science.abl4784 (2021).

3 Altarawneh, H. N. et al. Effects of Previous Infection and Vaccination on Symptomatic Omicron Infections. N Engl J Med, doi:10.1056/NEJMoa2203965 (2022).

4 Iketani, S. et al. Antibody evasion properties of SARS-CoV-2 Omicron sublineages. Nature 604, 553–556, doi:10.1038/s41586-022-04594-4 (2022).

5 Liu, L. et al. Striking antibody evasion manifested by the Omicron variant of SARS-CoV-2. Nature 602, 676–681, doi:10.1038/s41586-021-04388-0 (2022).

6 Planas, D. et al. Reduced sensitivity of SARS-CoV-2 variant Delta to antibody neutralization. Nature 596, 276–280, doi:10.1038/s41586-021-03777-9 (2021).

7 Wang, P. et al. Antibody resistance of SARS-CoV-2 variants B.1.351 and B.1.1.7. Nature 593, 130–135, doi:10.1038/s41586-021-03398-2 (2021).

8 Wang, Q. et al. Antibody evasion by SARS-CoV-2 Omicron subvariants BA.2.12.1, BA.4, and BA.5. bioRxiv, doi:https://doi.org/10.1101/2022.05.26.493517 (2022).

9 Gandhi, S. et al. De novo emergence of a remdesivir resistance mutation during treatment of persistent SARS-CoV-2 infection in an immunocompromised patient: a case report. Nat Commun 13, 1547, doi:10.1038/s41467-022-29104-y (2022).

10 Baden, L. R. et al. Efficacy and Safety of the mRNA-1273 SARS-CoV-2 Vaccine. N Engl J Med 384, 403–416, doi:10.1056/NEJMoa2035389 (2021).

11 Chen, P. et al. SARS-CoV-2 Neutralizing Antibody LY-CoV555 in Outpatients with Covid-19. N Engl J Med 384, 229–237, doi:10.1056/NEJMoa2029849 (2021).

12 Dougan, M. et al. Bebtelovimab, alone or together with bamlanivimab and etesevimab, as a broadly neutralizing monoclonal antibody treatment for mild to moderate, ambulatory COVID-19. medRxiv, doi:https://doi.org/10.1101/2022.03.10.22272100 (2022).

13 Gupta, A. et al. Early Treatment for Covid-19 with SARS-CoV-2 Neutralizing Antibody Sotrovimab. N Engl J Med 385, 1941–1950, doi:10.1056/NEJMoa2107934 (2021).

14 Polack, F. P. et al. Safety and Efficacy of the BNT162b2 mRNA Covid-19 Vaccine. N Engl J Med 383, 2603–2615, doi:10.1056/NEJMoa2034577 (2020).

15 Sadoff, J. et al. Safety and Efficacy of Single-Dose Ad26.COV2.S Vaccine against Covid-19. N Engl J Med 384, 2187–2201, doi:10.1056/NEJMoa2101544 (2021).

16 Weinreich, D. M. et al. REGN-COV2, a Neutralizing Antibody Cocktail, in Outpatients with Covid-19. N Engl J Med 384, 238–251, doi:10.1056/NEJMoa2035002 (2021).

17 Beigel, J. H. et al. Remdesivir for the Treatment of Covid-19 - Final Report. N Engl J Med 383, 1813–1826, doi:10.1056/NEJMoa2007764 (2020).

18 Gottlieb, R. L. et al. Early Remdesivir to Prevent Progression to Severe Covid-19 in Outpatients. N Engl J Med 386, 305–315, doi:10.1056/NEJMoa2116846 (2022).

19 Fischer, W. A., 2nd et al. A phase 2a clinical trial of molnupiravir in patients with COVID-19 shows accelerated SARS-CoV-2 RNA clearance and elimination of infectious virus. Sci Transl Med 14, eabl7430, doi:10.1126/scitranslmed.abl7430 (2022).

20 Jayk Bernal, A. et al. Molnupiravir for Oral Treatment of Covid-19 in Nonhospitalized Patients. N Engl J Med 386, 509–520, doi:10.1056/NEJMoa2116044 (2022).

21 Painter, W. P. et al. Human Safety, Tolerability, and Pharmacokinetics of Molnupiravir, a Novel Broad-Spectrum Oral Antiviral Agent with Activity Against SARS-CoV-2. Antimicrob Agents Chemother, doi:10.1128/AAC.02428-20 (2021).

22 Chan, A. P., Choi, Y. & Schork, N. J. Conserved Genomic Terminals of SARS-CoV-2 as Coevolving Functional Elements and Potential Therapeutic Targets. mSphere 5, doi:10.1128/mSphere.00754-20 (2020).

23 V’Kovski, P., Kratzel, A., Steiner, S., Stalder, H. & Thiel, V. Coronavirus biology and replication: implications for SARS-CoV-2. Nat Rev Microbiol 19, 155–170, doi:10.1038/s41579-020-00468-6 (2021).

24 Stevens, L. J. et al. Mutations in the SARS-CoV-2 RNA dependent RNA polymerase confer resistance to remdesivir by distinct mechanisms. Sci Transl Med, eabo0718, doi:10.1126/scitranslmed.abo0718 (2022).

25 Szemiel, A. M. et al. In vitro selection of Remdesivir resistance suggests evolutionary predictability of SARS-CoV-2. PLoS Pathog 17, e1009929, doi:10.1371/journal.ppat.1009929 (2021).

26 Hoffman, R. L. et al. Discovery of Ketone-Based Covalent Inhibitors of Coronavirus 3CL Proteases for the Potential Therapeutic Treatment of COVID-19. J Med Chem 63, 12725–12747, doi:10.1021/acs.jmedchem.0c01063 (2020).

27 Xie, X. et al. An Infectious cDNA Clone of SARS-CoV-2. Cell Host Microbe 27, 841–848 e843, doi:10.1016/j.chom.2020.04.004 (2020).

28 Unoh, Y. et al. Discovery of S-217622, a Noncovalent Oral SARS-CoV-2 3CL Protease Inhibitor Clinical Candidate for Treating COVID-19. J Med Chem 65, 6499–6512, doi:10.1021/acs.jmedchem.2c00117 (2022).

29 Mukae, H. et al. A Randomized Phase 2/3 Study of Ensitrelvir, a Novel Oral SARS-CoV-2 3C-like Protease Inhibitor, in Japanese Patients With Mild-to-Moderate COVID-19 or Asymptomatic SARS-CoV-2 Infection: Results of the Phase 2a Part. medRxiv, doi:https://doi.org/10.1101/2022.05.17.22275027 (2022).

30 Ferreira, J. C., Fadl, S. & Rabeh, W. M. Key dimer interface residues impact the catalytic activity of 3CLpro, the main protease of SARS-CoV-2. J Biol Chem 298, 102023, doi:10.1016/j.jbc.2022.102023 (2022).

31 Shu, Y. & McCauley, J. GISAID: Global initiative on sharing all influenza data - from vision to reality. Euro Surveill 22, doi:10.2807/1560-7917.ES.2017.22.13.30494 (2017).

32 (ASPR), H. O. o. t. A. S. f. P. a. R. COVID-19 Therapeutics Thresholds, Orders, and Replenishment by Jurisdiction, <https://aspr.hhs.gov/COVID-19/Therapeutics/orders/Pages/default.aspx> (2022).

33 Iketani, S. et al. The Functional Landscape of SARS-CoV-2 3CL Protease. bioRxiv, doi:https://doi.org/10.1101/2022.06.23.497404 (2022).

34 Jochmans, D. et al. The substitutions L50F, E166A and L167F in SARS-CoV-2 3CLpro are selected by a protease inhibitor in vitro and confer resistance to nirmatrelvir. bioRxiv, doi:https://doi.org/10.1101/2022.06.07.495116 (2022).

35 Zhou, Y. et al. Nirmatrelvir Resistant SARS-CoV-2 Variants with High Fitness in Vitro. bioRxiv, doi:https://doi.org/10.1101/2022.06.06.494921 (2022).

36 Xue, X. et al. Structures of two coronavirus main proteases: implications for substrate binding and antiviral drug design. J Virol 82, 2515–2527, doi:10.1128/JVI.02114-07 (2008).

37 Yang, S. et al. Synthesis, crystal structure, structure-activity relationships, and antiviral activity of a potent SARS coronavirus 3CL protease inhibitor. J Med Chem 49, 4971–4980, doi:10.1021/jm0603926 (2006).

38 Zhao, Y. et al. Crystal structure of SARS-CoV-2 main protease in complex with protease inhibitor PF-07321332. Protein Cell 13, 689–693, doi:10.1007/s13238-021-00883-2 (2022).

39 Sacco, M. D. et al. The P132H mutation in the main protease of Omicron SARS-CoV-2 decreases thermal stability without compromising catalysis or small-molecule drug inhibition. Cell Res 32, 498–500, doi:10.1038/s41422-022-00640-y (2022).

40 Liu, H. et al. Development of optimized drug-like small molecule inhibitors of the SARS-CoV-2 3CL protease for treatment of COVID-19. Nat Commun 13, 1891, doi:10.1038/s41467-022-29413-2 (2022).

41 Liu, L. et al. Potent neutralizing antibodies against multiple epitopes on SARS-CoV-2 spike. Nature 584, 450–456, doi:10.1038/s41586-020-2571-7 (2020).

42 Freed, N. E., Vlkova, M., Faisal, M. B. & Silander, O. K. Rapid and inexpensive whole-genome sequencing of SARS-CoV-2 using 1200 bp tiled amplicons and Oxford Nanopore Rapid Barcoding. Biol Methods Protoc 5, bpaa014, doi:10.1093/biomethods/bpaa014 (2020).

43 Annavajhala, M. K. et al. Emergence and expansion of SARS-CoV-2 B.1.526 after identification in New York. Nature 597, 703–708, doi:10.1038/s41586-021-03908-2 (2021).

44 Reed, L. J. & Muench, H. A SIMPLE METHOD OF ESTIMATING FIFTY PER CENT ENDPOINTS. American Journal of Epidemiology 27, 493–497, doi:https://doi.org/10.1093/oxfordjournals.aje.a118408 (1938).

45 Stirling, D. R. et al. CellProfiler 4: improvements in speed, utility and usability. BMC Bioinformatics 22, 433, doi:10.1186/s12859-021-04344-9 (2021).

46 Posada-Cespedes, S. et al. V-pipe: a computational pipeline for assessing viral genetic diversity from high-throughput data. Bioinformatics, doi:10.1093/bioinformatics/btab015 (2021).

47 Virtanen, P. et al. SciPy 1.0: fundamental algorithms for scientific computing in Python. Nat Methods 17, 261–272, doi:10.1038/s41592-019-0686-2 (2020).

48 Waskom, M. L. seaborn: statistical data visualization. Journal of Open Source Software 6, 3021 (2021).

49 Ye, C. et al. Rescue of SARS-CoV-2 from a Single Bacterial Artificial Chromosome. mBio 11, doi:10.1128/mBio.02168-20 (2020).

50 Chen, A. T., Altschuler, K., Zhan, S. H., Chan, Y. A. & Deverman, B. E. COVID-19 CG enables SARS-CoV-2 mutation and lineage tracking by locations and dates of interest. Elife 10, doi:10.7554/eLife.63409 (2021).

